# High-Order Organization of Guanine-Based Reflectors Underlies the Dual Functionality of the Fish Iris

**DOI:** 10.1101/240028

**Authors:** Dvir Gur, Jan-David Nicolas, Vlad Brumfeld, Omri Bar-Elli, Dan Oron, Gil Levkowitz

**Affiliations:** Department of Physics of Complex Systems, Weizmann Institute of Science, Rehovot, 7610001, Israel.; Department of Molecular Cell Biology, Weizmann Institute of Science, Rehovot, 7610001, Israel.; Institut für Röntgenphysik, Göttingen, 37077, Germany.; Department of Chemical Research Support, Weizmann Institute of Science, Rehovot, 7610001, Israel.

## Abstract

Many marine organisms have evolved a reflective iris to prevent unfocused light from reaching the retina. The fish iris has a dual function, both to camouflage the eye and serving as a light barrier. Yet, the mechanism that enables this dual functionality and the benefits of using a reflective iris have remained unclear. Using synchrotron micro-focused diffraction, cryo-SEM imaging and optical analyses on zebrafish at different stages of development, we show that the complex optical response of the iris is facilitated by the development a high-order organization of multilayered guanine-based crystal reflectors and pigments. We further demonstrate how the efficient light reflector is established during development to allow the optical functionality of the eye, already at early developmental stages. These results shed light on the evolutionary drive for developing a compact reflective iris, which is widely used by many animal species.

**Significance Statement:** The fish iris is an exquisite example of nature’s remarkable engineering where specialized cells, dubbed iridophores, produce an efficient light reflector made of guanine-based crystals. This unique structure of the fish iris serves a dual function: In addition to its role as a light barrier, the iris has a second role of camouflaging the eye by creating a silvery reflectance, which merges with the fish skin. The underlying mechanism that enables the aforementioned dual functionality of the fish iris as well as the structural morphogenesis of the guanine reflector during embryonic development, remained unclear. We show that complex optical response of the iris is facilitated by the establishment of a high-order organization of multilayered guanine-based crystal reflectors and pigments.

## Introduction

In the eyes of numerous organisms, including mammals, reptiles, birds and fish, light is absorbed by photoreceptors in the dorsally located retina. In order to obtain a high-contrast image, only light that passes through the pupil and is focused by the lens should reach the retina. Hence, a key function of the iris is to prevent the passage of unfocused light. Irises can be classified either as absorptive, or as reflective or scattering, according to their operating principle with regard to incident light ^1,2^. Many irises of the absorptive type, including in the human eye, contain melanin pigments that efficiently absorb a broad spectrum of visible light ^1,3^. An alternative strategy is employed by numerous marine organisms, in which the outer surface of the iris is highly reflective ^2,4-8^.

The reflecting iris of many fish species consists of intracellular crystals that are formed by the nucleobase guanine ^9-12^. These guanine crystals are assembled in specialized cells, named iridophores, which are located mainly in the eyes and skin ^2,13-15^. In addition to its function as a light barrier, the iris of fish has a second role of camouflaging the eye by creating a silvery reflection similar to that of fish skin ^2,9,11^. Both these functions require the iris to reflect light efficiently. However, for effective camouflage the iris should reflect light in a highly directional manner ^9,11^, whereas to block light the iris should reflect or scatter light independently of angle or wavelength ^1,3^.

Although the dual function of the evolutionarily conserved fish iris has long been known ^2,11^, the underlying mechanism that enables this duality has remained unclear. Here, we unravel the unique ultrastructure of iridophores in the zebrafish iris, which comprises multilayered organization of the guanine-based crystals. We directly associate the iris crystal organization to simulated and measured reflectance spectrum. We further show that the structural organization of the developing zebrafish iris is ideally designed to reflect light efficiently at a very early stage of development, which correlates well with the reported high visual acuity of the larva. Mapping of crystal orientations across the entire iris revealed the high-order crystal organization, which enables the dual function of the iris.

## Results

### Zebrafish iris is a multilayered optical system

To elucidate iris anatomy and the subcellular organization of the guanine-based reflectors, we used the zebrafish eye as a model system. In it, the lens is surrounded by a silvery iris, termed *argenta*, which is exposed to direct external light (Fig. 1, A to C). We first imaged the entire zebrafish eye using micro-CT (Fig. 1, D to F), in which the guanine-based crystal layers of the iris are clearly visible due to their high density. Serial scans showed that the zebrafish iris extends to the dorsal part of the eyeball, where it forms a reflecting mirror behind the retina, termed the *tapetum lucidum* (Fig. 1, B, C and E). The upper, outermost layer of the eye was devoid of the reflecting guanine-based layer and, instead, was covered by melanin pigments (Fig. 1B). In tomographic images of sagittal slices through the dorsal (upper) or ventral (lower) part of the eye, the iris surface appeared as a continuous layer across the entire section (Fig. 1E), whereas in a sagittal slice through the center of the eye, the two sides of the iris were separated by the lens (Fig. 1F).

**Fig. 1.**
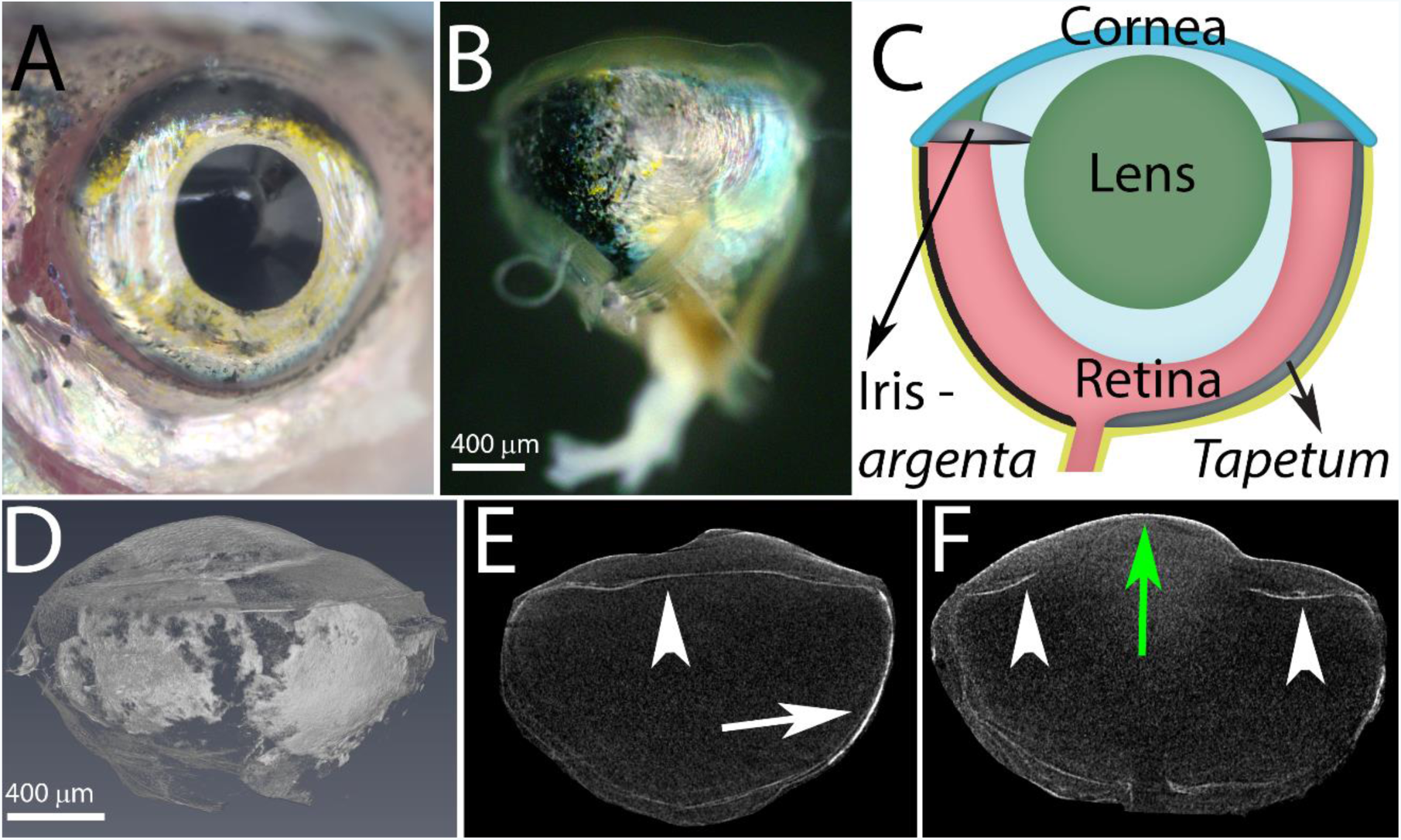
The anatomy of the zebrafish eye. A) A top view of the zebrafish eye showing the silvery iris. B) A side view of the fish eyeball showing the *tapetum lucidum*, a silvery layer, located at the bottom part of the eyeball (right) and a pigmented layer visible at the upper part of the eyeball (left). C) A schematic representation of the eye showing the cornea (blue), lens (green), and retina (light red), as well as the guanine-based reflecting layers, the iris *argenta* and the *tapetum lucidum* (silver) and the pigmented layer (black). D-F) Micro-CT images of the zebrafish eye: a volume rendering of the whole eye (D), a sagittal view of the lower part of the eye (E), and a sagittal view of the center of the eye (F). Arrowheads indicate the iris *argentum*, white arrow indicates the *tapetum lucidum* and green arrow indicates the cornea. Scale bars: 400 μm.

To visualize the organization of the guanine-based reflectors in the adult zebrafish iris under near-physiological conditions we used cryo-SEM, which allows preservation of intact guanine crystals in their native cellular context (Fig. 2 A and B). Imaging of coronal sections through the eye revealed that the iris is composed of three different layers: an upper layer of ordered iridophores, a middle layer of disordered iridophores, and a thin layer (~5 μm) of pigment-forming melanophores, which lies beneath the two iridophore layers (Fig. 2, A and E). The cryo-SEM analysis also showed that the outermost, ordered iridophore layer was composed of monodisperse elongated hexagonal guanine crystals, which were stacked on top of each other (Fig. 2, A and C). These crystals displayed an average thickness of 25±7 nm (*n*=150) and an average cytoplasm spacing of 142±25 nm (*n*=140). In the adjacent disordered layer, the spacing between crystals was highly variable and their arrangement appeared random, with no preferred orientation (Fig. 2, A and D). Notably, the disordered iridophore layer was present only in the iris *argenta* and was missing from the *tapetum lucidum*.

**Fig. 2.**
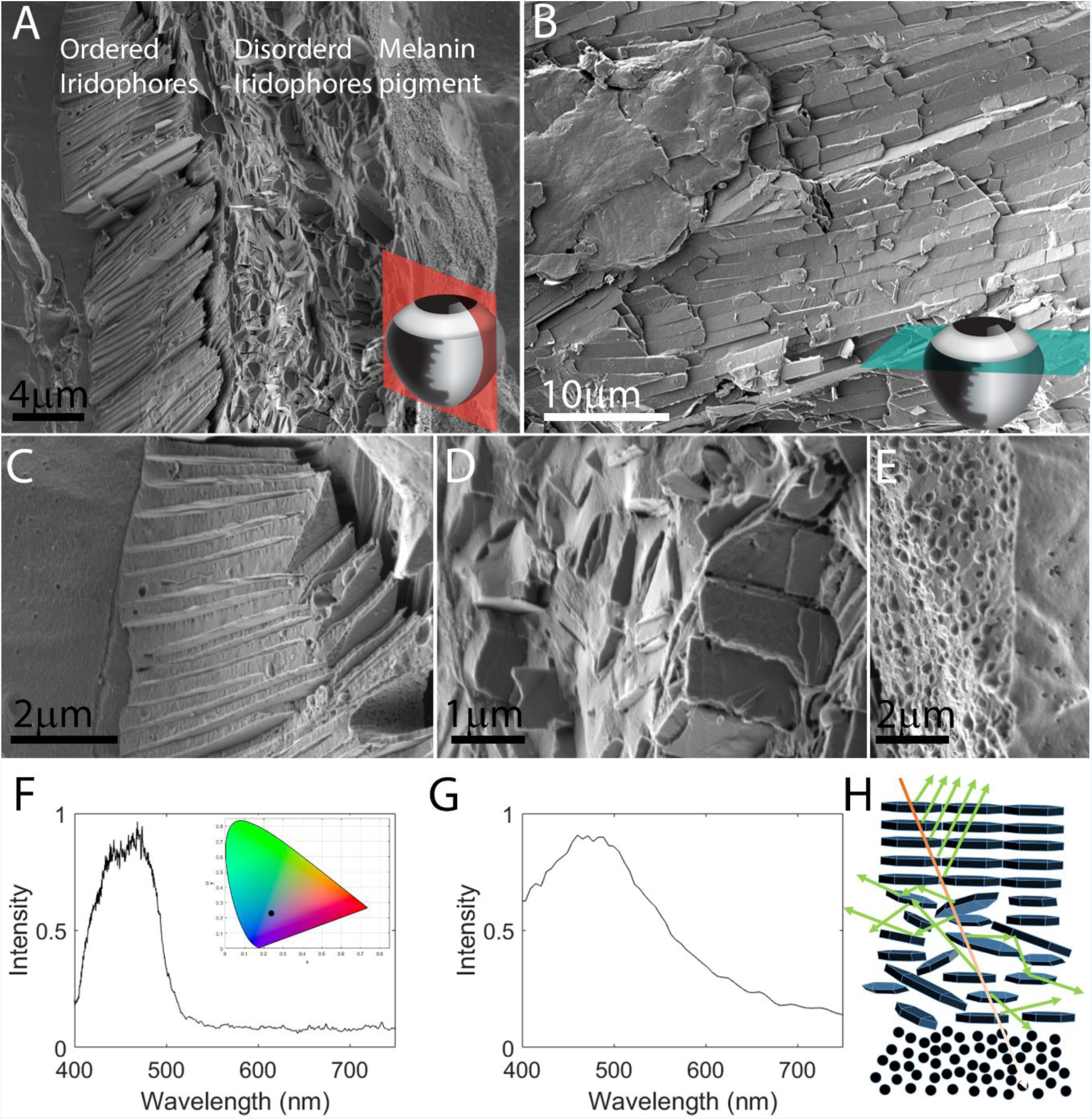
The ultrastructure and optical properties of the iris. A) A coronal section through the iris showing three distinct layers: (i) ordered iridophores, (ii) disordered iridophores, and (iii) a pigmented layer. The section was taken from the periphery of the eye where the iris is curved and, thus, the crystals in the ordered layer are tilted. B) A top view of the iris obtained by a transverse section through the eye shows the elongated guanine crystals packed one next to the other, perfectly tiling the surface of the iris. The anatomical location of sections are illustrated in the bottom-right corners in A,B. C-E) Show higher magnifications of the different iris layers: ordered iridophores (C), disordered iridophores (D), and pigmented layer (E). A-E) Show Cryo-SEM micrographs of high-pressure frozen, freeze-fractured iris F) Graph showing measured reflectance from the iris, revealing a broad peak centered around 450 nm. Inset shows the color of reflected light (black dot) plotted on a 1931 CIE chromaticity space diagram. G) Simulated reflectance of the ordered layer calculated based on the data in high-resolution cryo-SEM micrographs H) A sketch illustrating the different paths of light traveling through the multi-layered iris. Scale bars: 4 μm (A), 10 μm (B), 2 μm (C,E), 1μm (D).

We next imaged a horizontal section of the eye (Fig. 2B), where the ordered layer was seen as an array of tightly packed crystals perfectly tiling the surface of the iris. The tiling of the crystals was extremely compact, to the extent that it was difficult to separate between iridophores. These results show that in the iris of zebrafish eye, the *argenta* and *tapetum lucidum* form a continuous layer. Yet, whereas the *tapetum* is composed only from one ordered layer of iridophores, the *argenta* has a second, disordered guanine-based layer underlaid by a thin melanosome-based pigment layer.

### The ordered iridophore layer reflects most of the blue-green light

We hypothesized that the three-layer composition of the iris underlies a complex optical response, which combines wavelength-dependent light reflection, scattering and absorption. Because impinging light will first hit the reflectors of the outermost layer of the iris, we assumed that the majority of light would be reflected from the ordered layer. To simulate the reflectance of the ordered layer, we analyzed our cryo-SEM data, namely the spacing between crystals, using the Monte Carlo transfer matrix method (see Materials and Methods). The analysis showed that the ordered layer mostly reflects blue-green light, which peaked at 450 nm (Fig. 2G). To correlate our simulated results with actual optical measurements, we used a custom-built light microscope equipped with a spectrophotometer. Indeed, spectral measurements showed that approximately 80-90% of the blue-green light was reflected from the iris (Fig. 2F). The agreement between simulated calculations and optical measurements suggests that the majority of the blue-green light is indeed reflected from the ordered iridophore layer of the zebrafish iris.

Based on the combined results of ultrastructure imaging, light reflectance simulations and real-time optical measurements, we propose a model for light traveling through the iris (Fig. 2H). The ordered iridescent layer reflects approximately 80-90% of the impinging blue-green light. The residual light that passeses through the first layer is scattered by the disordered layer. Finally, the thin pigmented layer likely absorbs any light passing the two guanine-based layers, including light that is forward-scattered from the disordered layer. In addition, the scattering of light by the disordered layer, as it likely introduce an angular spread to the incoming light, increases the probability for light to be absorbed by the melanin pigment.

### The zebrafish iris is functional at early developmental stages

Zebrafish reproduce by external fertilization, which means that the offspring must be self-sufficient very early in life. High visual acuity is pertinent to the survival of free-swimming zebrafish, which rely on visual stimuli for prey capture ^16-18^. Previous studies showed that iridophores in zebrafish iris could be detected as soon as 3dpf ^7^. We therefore predicted that the zebrafish iris would be highly functional at early developmental stages. To test this hypothesis, we analyzed the optical properties, iridophore ultrastructure and crystal organization of the zebrafish iris at different stages of development. Light microscopy revealed that iridophores were indeed clearly visible in the iris already at 3 days post fertilization (dpf), where they intially occupy the inner circumferential layer, lining the inner radius of the iris around the lens (Fig. S1). At 7 dpf, iridophores covered only about half of the iris surface (Fig. 3A). At 14 dpf, the iridophores occupied a larger area of the iris; however, large dark patches showing the underling pigmented layer, were still visible. By 21 and 28 dpf, the iridophores covered nearly the entire surface of the iris (Fig. 3, C and D).

**Fig. 3.**
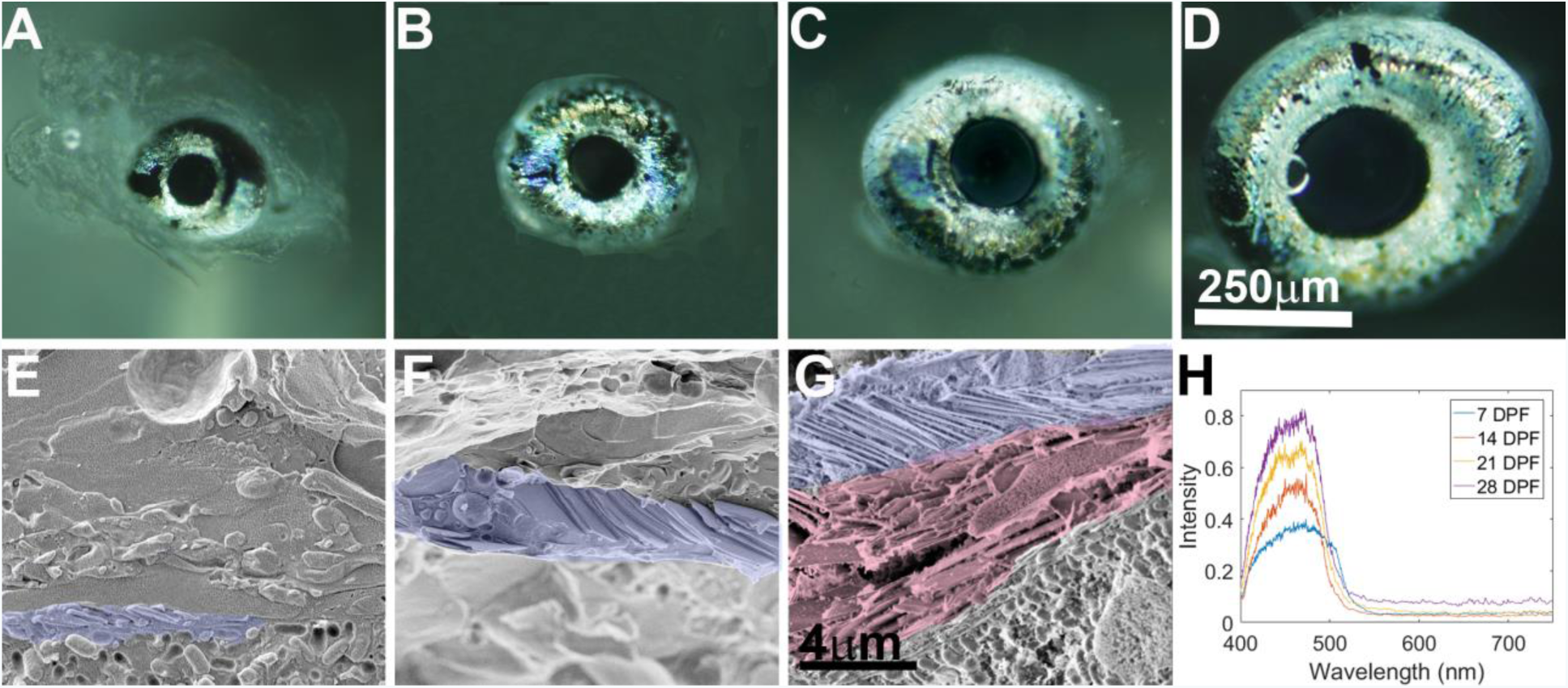
Iris development. Iris morphology, ultrastructural and optical properties were examined during development at 7 (A,E), 14 (B,F), 21 (C,G) and 28 (D) dpf. A-D) Light microscope images of the larva iris at different stages. E-G) Cryo-SEM micrographs of high-pressure frozen, freeze-fractured larva iris show the ultrastructure of the developing iris. Whereas at 7 and 14 dpf only the ordered layer is seen, by 21 dpf a second, disordered layer starts to appear. Ordered iridophores (purple) and disordered iridophores (light red) are shown in pseudo-colors H) Graph showing measured reflectance of irises from 7, 14, 21 and 28 dpf larvae, revealing a peak reflectance centered around 450 nm. Scale bars: 250 μm (A-D), 4 Ϝm (E-G).

To elucidate the development of the iris ultrastructure, we performed cryo-SEM imaging at corresponding development stages. We found that at 7 dpf, the guanine crystals in the clearly visible iridophores were arranged in relatively ordered parallel stacks, similar to the adult iris (Fig. 3E). However, at this stage, only one layer of iridophores was visible. Moreover, individual crystals were smaller than those observed in adults and, in certain cases, the membranes engulfing the crystal seemed less tight than in adults (Fig. 3E). At 14 dpf, still only one layer of iridophores was present, but the crystal stacks were more ordered compared to 7 dpf (Fig. 3F). By 21 dpf, iris thickness increased considerably and the second layer of iridophores was sometimes visible(Fig. 3G), usually in the areas closest to the lens (Fig. S2). At this age, guanine crystals and iridophore ultrastructure resembled those of adult fish, suggesting that the iris is already fully functional as a reflecting light barrier.

To determine the functionality of the iris during development, we measured the reflectance spectrum at corresponding stages (Fig. 3H). Despite the incomplete crystal layers and iridophore coverage at larval stages, a wide reflectance peak centered on 450 nm wavelength was measured from a very early stage (7 dpf) until adulthood (Fig. 3H). The reflectance intensity increased with age, averaging at ~40% for 7 dpf larva and reaching ~80% by 28 dpf, very close to the levels observed in adults. Taken together, these results indicate that the larval iris functions as an efficient light-barrier already at 7 dpf, and that this initial functionality is preserved and optimized throughout eye development. These findings are consistent with the reported high visual acuity of very early stage zebrafish larvae ^16-18^.

### Synchrotron-based diffraction imaging of the whole iris reveals high order organization

Having found that light is reflected mostly from the outermost ordered iridophore layer, we hypothesized the existence of an intrinsic, high-order organization of crystal stacks within this layer. To examine this, we sought to map the orientation of the guanine crystals stacks across the entire iris. Because the adult zebrafish iris is composed of millions of crystals extending across several square millimeters, mapping crystal orientations accurately using high-resolution imaging techniques such as cryo-SEM was not feasible, whereas modalities for imaging relatively large areas, such as micro-CT, would not provide sufficient resolution to determine the orientation of individual stacks. Therefore, we opted to use scanning x-ray diffraction combined with a micro-focused synchrotron beam. Thus, the x-ray beam continuously scanned across the sample and the diffraction patterns were collected with a single-photon counting flat panel detector at regular time intervals. The scanning speed was set such that a diffraction pattern was collected every 4 μm. Thus, a single scan could contain over 400,000 diffraction patterns, covering the entire surface area of the iris at an overall scan time of about 1 hour per sample. To analyze the data in a standardized, robust and model-independent fashion, we used a custom Matlab-based principal component analysis (PCA) based on Bernhardt et al. ^19^ as part of a Nano-diffraction toolbox Nicolas et al ^20^ (Materials and Methods, supporting information). Using this PCA algorithm, we extracted the anisotropy and orientation of light scattering and, subsequently, inferred the orientation of the crystals.

Based on our TEM studies (Fig. S3), the crystals in the fish iris are anhydrous β-guanine (100) crystal plates, (Fig. 4C). Thus, the (100) plane is in diffraction for crystals that are oriented almost edge-on to the beam (see Fig. 1A color bar for crystal orientation). Mapping the anisotropy and orientation of the (100) diffraction plane revealed two regions of crystal stacks, an inner layer that surrounded the lens and an outer layer surrounding the outer radius of the eye (Fig. 4A). The crystals in the two layers were positioned tangent to the iris circumference, lining the top section of the eye. Remarkably, the orientation of the crystal stacks at the inner and outer radii followed the curvature of the iris (Fig. 4A). In the outer layer, the crystals extended up to the point where the eye is protruding out from the fish head, suggesting that they prevent light that impinges upon the eye from the side from distorting the fish vision (Fig. 4F and Fig. S4). Accordingly, the crystals at the inner layer, which surrounds the lens, would reflect light that is impinged upon the lens sideways, and thus limit the angular acceptance of the eye, leading to a reduction in spherical aberrations. This inner circumference layer may also provide a tighter seal for light at the interface between the lens and the iris (Fig. 4F).

**Fig. 4.**
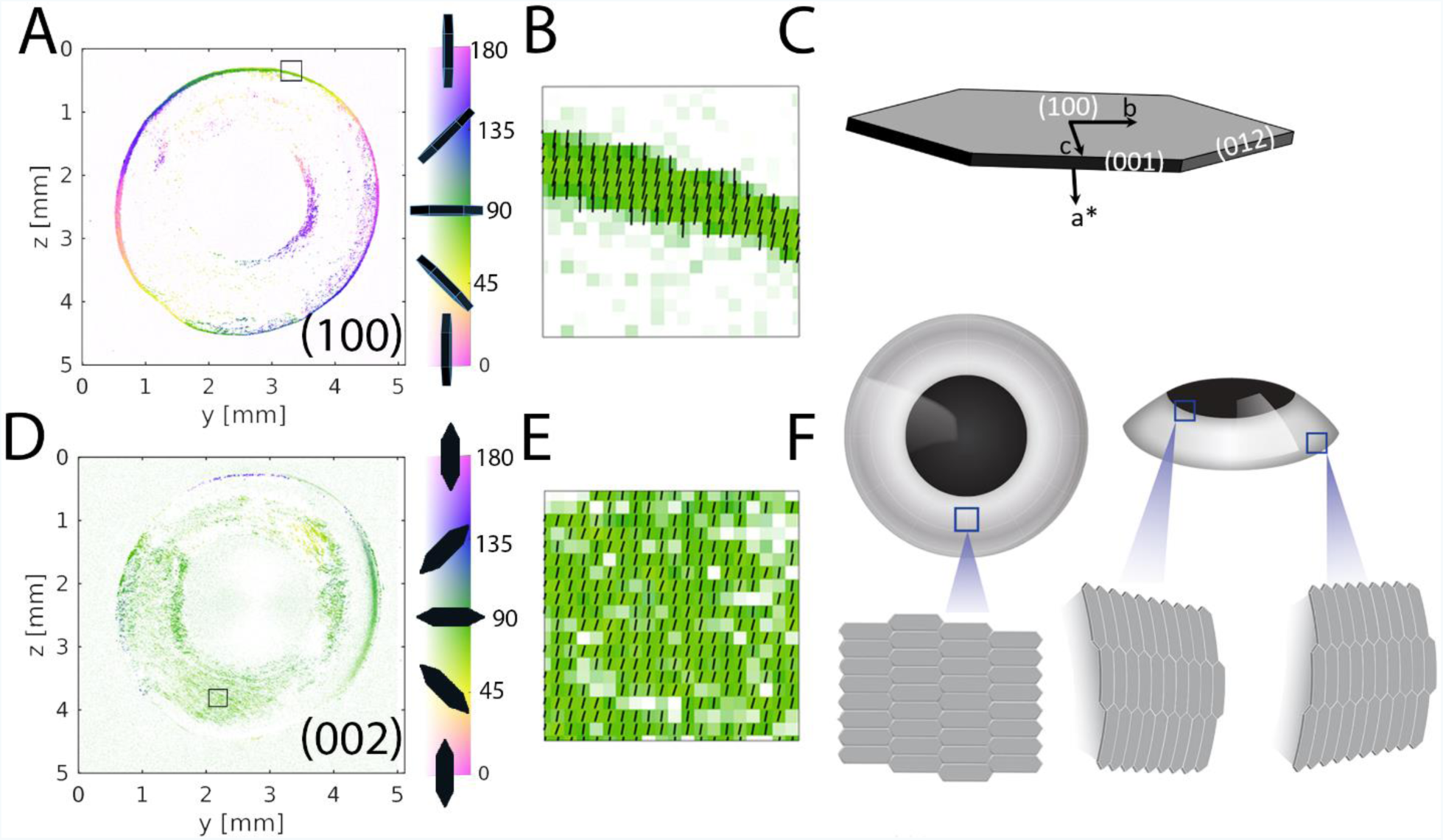
Synchrotron-based diffraction imaging. The orientation of the different crystal planes was mapped across the entire iris using scanning x-ray diffraction combined with a micro-focused synchrotron beam. A,D) Maps showing the orientations of the (100) and (002) guanine crystal diffraction planes, respectively. The color gradient (right bar) from white to saturated color corresponds to the respective low and high anisotropy levels. B,E) Magnifications of the squared regions marked in (A) and (D), respectively. C) Schematic illustration of fish guanine crystal, showing the relations between crystal morphology and the crystallographic axes and faces. F) Schematic illustration showing the orientations of the guanine plate-like crystals at different regions of the eye.

The perpendicular (002) plane should only be in diffraction for crystals that are positioned almost perpendicular to the beam (Fig. 4C). Mapping the (002) diffraction plane across the iris showed that the orientation of most of the crystals was parallel to the iris surface (Fig. 4D), suggesting that they would efficiently reflect light impinging on the surface (Fig. 4F). Due to the curvature of the iris close to its periphery, the crystal were tilted and were thus out of diffraction. Nonetheless, these regions are still tiled by crystals, as observed by mapping the diffraction of all the different crystal planes, regardless of their orientation (Fig. S5B).

Remarkably, the crystals that tile the surface of the iris were aligned throughout the entire iris, covering several square millimeters in adult fish. This co-orientation of crystals across tens of thousands of cells suggests that the concerted organization of crystals is highly regulated. Presumably, cell communication between iridophores facilitates orienting the crystals across the iris in a synchronized manner. Such coordinated cell communication mechanism was previously reported for other types of pigment cells during zebrafish color pattern formation ^21-24^. Light reflection from a flat layer of co-orientated crystals could be analogous to the specular reflection from a flat mirror, which is characterized by a high degree of directionality. Indeed, Fourier transform reflection analysis indicated that reflection from the iris is highly directional (Fig. S6).

## Discussion

In zebrafish, the eyes are located on both sides of the head such that the iris surfaces are at an oblique angle to impinging sunlight. Thus, sunlight should be reflected from the iris at an obtuse angle, preserving the down welling direction of the light. This may be beneficial for camouflage in shallow waters, where upwelling iridescence is relatively minute ^25^. Thus, light reflected or scattered in any direction other than downwelling would be highly conspicuous, revealing the location of the fish to potential predators. Furthermore, the co-orientation of the crystals may allow perfect surface tiling and, thus, minimize surface defects. Such defects may lead to transmission of light through the iris and onto the retina, reducing the contrast of the obtained image.

Having both an ordered and a disordered layer enables the dual functionality of the iris, an efficient light barrier that also provides camouflage to the normally black uveal tract. The first ordered layer is an efficient directional light reflector and, at most angles, incident light will be reflected efficiently. However, at very obtuse angles, at which light is almost parallel to the reflectors (for example, sunlight scattered by an object in the water), or when light is at the red edge of the visible spectrum, the efficiency of the reflectors is reduced dramatically. Having a second layer of varying crystal orientations could provide a solution to this problem. A second functional benefit for having two iris layers relates to the spectral environment of the eye. In the marine environment, the longer wavelengths are rapidly absorbed in the water, leaving mostly blue-green light ^25,26^. Furthermore, the opsins in the zebrafish retina are much more sensitive to shorter wavelengths than to longer ones ^18,27,28^. Thus, the iris should be very efficient in reflecting shorter wavelength, while still being able to reflect lower levels of longer wavelengths. The two superimposed layers in the iris provide an elegant solution to these requirements. The ordered layer has a peak reflectance for shorter wavelengths and the disordered layer scatters most wavelengths, but to a lesser extent. The organization of the plate-like crystals in the ordered layer changes considerably in different regions of the iris. The crystals in the center of the iris are parallel to the iris surface, whereas at the edges they are perpendicular to the surface. This high-order organization allows the iris to efficiently block laterally impinging light and reduces spherical aberration. As mentioned, the co-orientation of the crystals allows for flawless tiling across the iris surface, thereby providing a tight seal for light. Interestingly, in certain albino fish the peripheral rods evade light damage although the inner iris epithelium lacks melanin, suggesting that the iris *argenta* also provides sufficient light screening to protect these photoreceptors ^29^.

Our analyses of the developing iris may explain the evolutionary choice of a reflective rather than absorptive iris in fish. The diameter of the 5 dpf larval eye is approximately 150 μm, increasing up to ~350 μm at 21 dpf. Taking into account the volume of the other eye parts, the space that the iris may occupy is very limited. An ordered reflecting layer is compact and, thus, advantageous under these conditions. Indeed, we found that the ordered reflecting layer, which is only ~3 μm thick, provides about 60% reflection of the blue-green light. In comparison, a melanin-based iris would require a layer of about 15 μm in thickness to absorb a similar amount of light ^30^.

Lastly, the ordered crystal layer, which mostly reflects blue-green light, is advantageous because these wavelengths are the most abundant in the marine environment. The increase in iris reflectivity with age is mostly due to increase in the coverage area by iridophores, but probably also due to the thickening of the ordered reflecting layer. The thin layer of ordered reflectors blocks a sufficiently broad spectrum of light and should allow the developing larvae to form an image, albeit of low contrast, at a very early age. The second disordered layer, which develops later, scatters a broader spectrum of light and, thereby, enables the formation of clearer images of higher contrast. This developmental strategy enables the eye to become functional at a very early stage and remain functional throughout development, as iris efficiency increases.

The fish iris is an exquisite example of nature’s remarkable engineering, where the specialized iridophores produce an efficient light reflector made of guanine-based crystals. We provide a biophysical mechanism for the dual functionality of the iris, an efficient light barrier and a camouflage for the fish eye. The complex optical response of the iris is achieved by a combination of wavelength-dependent light reflection from the ordered layer, scattering by the disordered layer, and absorption by the pigmented layer. The high-order organization of the reflecting layer both optimizes the light reflection properties of the iris, stops laterally impinging light, and provides the eye with camouflage. From an evolutionary-developmental perspective, the drive for developing a reflective iris could have been the small size of the eye during early development, which favors the more compact reflecting iris over the absorbing one for preventing light from reaching the retina.

## Methods

### Zebrafish lines and maintenance

Zebrafish were raised and bred according to the Weizmann Institute Animal Care and Use Committee (IACUC). The zebrafish lines used were either AB or TL wild types.

### Chromatic imaging

High-quality color images were collected using an inverted microscope (Eclipse Ti-U, Nikon) equipped with a Nikon high-definition color camera head (DS-Fi2, Nikon) or using a Zeiss Axioplan microscope (Zeiss, Jena, Germany).

### Cryo-scanning electron microscopy (Cryo-SEM)

For adult zebrafish, fixed eyes were embedded in 3% agar using PBS as the medium. Coronal or transverse sections were cut at a thickness of 200 μm using a vibratome (VT1000-S, Leica). For zebrafish larvae, either fixed or fresh sacrificed larvae (5-28 dpf) were used. The sections or larvae were then sandwiched between two metal discs (3 mm diameter, 0.1 mm cavities) and cryo-immobilized in a high-pressure freezing device (HPM10; Bal-Tec). The frozen samples were mounted on a holder under liquid nitrogen and transferred to a freeze-fracture device (BAF60; Bal-Tec) using a vacuum cryo-transfer device (VCT 100; Bal-Tec), where they were coated with a 4-nm-thick layer of Pt/C. Samples were then observed by high-resolution SEM (Ultra 55, Zeiss) using secondary electron/backscattered electron and an in-lens detector, maintaining the frozen-hydrated state by using a cryo-stage operating at a working temperature of −120°C. Measurements of crystal thickness and cytoplasm spacing were taken from the cryo-SEM micrographs.

### X-ray micro-CT

Micro-CT scans were performed using a Micro XCT-400 (Zeiss X-ray Microscopy, California, USA). Whole zebrafish eyes, including the optic nerve, were placed in a plastic pipette tip, which had been sealed by melting using a flame. To prevent dehydration of the sample, the tips were partially filled with water and the eye was held above the water in a saturated water vapor atmosphere. X-ray micro-CT measurements were performed on a total of 5 whole, hydrated zebrafish eyes. The tomographic volumes were obtained by taking 1100 projections over 180° at 30 KV and 150 μA, and the isotropic voxel size in the reconstructed volume was 2.2 μm.

### Reflectivity measurements

The reflectivity of the zebrafish iris was measured on either fixed or recently removed fresh eyes, which were immersed in PBS and placed underneath a cover slip held in place with silicon grease. The eyes were positioned such that the iris was facing directly upwards. Impinging light was approximately normal to the objective lens. Reflectivity was measured as described in detail in ^14^ with the exception of using a Shamrock 303i spectrometer (Andor) equipped with an iDus 420 CCD camera and cooled to −70°C. Briefly, we used a custom-built microscope consisting of a microspectrophotometer, two CCD cameras, and a high numerical aperture objective, which enabled imaging the iris while obtaining both the reflectance spectrum and the Fourier transform of the reflectance for the same location in the sample. The light source was a halogen lamp coupled to an optical fiber, which guided the light into the microscope.

Imaging was first used to determine the correct focal point for the light source. Then the reflectance spectrum was used to determine the reflectance intensity, which was normalized to the reflectance of a silver mirror. Light was then imaged through a beam splitter onto the back aperture of an objective (Olympus, UPLSAPO 60XW, NA 1.2). The objective was used both to illuminate a wide area (~250 μm in width) and to collect the scattered light. The collected light was directed to one of three different paths by a set of folding mirrors. In the first path, the sample was imaged onto a CCD camera (Mintron, MTV 13 V5Hc). In the second path, the Fourier transform of the scattered light was captured by imaging the back aperture of the objective onto a similar CCD camera. In the third path, the light was collected and coupled into a fiber, which guided the light into a spectrometer. The sample was placed on top of translational stage and goniometer, such that both its position and orientation could be controlled.

### Reflectivity simulations

The reflectivity spectrum was simulated based on crystal thicknesses and spacing obtained from cryo-SEM images using a Monte Carlo transfer matrix calculation, as described in detail in the supporting information of ^31^. In brief, the percentage of reflectivity was calculated by averaging 500 runs, assuming normal incident light. Each layer was characterized by two variables: n_j_, a refractive index, and d_j_, which is the layer thickness randomly picked from the experimental distribution. Thus, for each layer we defined the following 2 × 2 matrix:

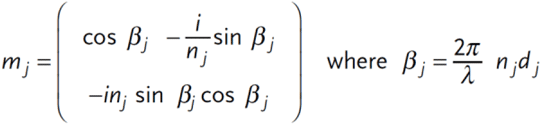

The set of k double layers was characterized by an overall reflectivity 2 × 2 matrix:

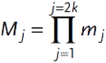

The reflectivity was extracted from the following equation:

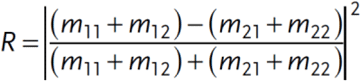

The refractive index of the guanine crystal plates was taken as 1.83, which was the refractive index in the direction of the impinging light. The weak dependence of the refractive index on wavelength was neglected, assuming that all the interfaces, i.e., inside a crystal stack and between stacks, were parallel. We also assumed no correlation between the crystal spacings within a single crystal stack.

### Transmission Electron Microscopy (TEM) imaging

To extract the guanine crystals, eyes were embedded in 7% agar and cut into 100 μm horizontal sections using a vibratome. The sections were then homogenized and the crystals were concentrated using centrifugeation. A suspension of the crystals in DDW was then removed and a drop was applied to a glow-discharged carbon-coated, copper TEM grid. The suspension was allowed to settle for 30 seconds and were then blotted. The TEM grids were observed using an FEI Tecnai T12 TEM operated at 120 kV. Images and diffraction patterns were recorded on a Gatan OneView camera using imaging and diffraction modes respectively. The observed electron diffraction patterns of the crystals (in set in Fig. S5) correspond to anhydrous β-guanine^32^.

### Analysis of Diffraction Data

Data correction steps were necessary, before scattering data could be analyzed. First, invalid detector pixels were masked such that their respective value were not taken into account in the analysis. Secondly, the absorption of a semi-transparent capillary used as a beamstop holder was corrected for by pixel wise multiplication with a correction matrix. Thirdly, background was subtracted from each scattering pattern to base the PCA analysis solely on the scattered intensity due to crystalline reflections. Lastly, to reduce data load and to improve the speed of calculation, the PCA analysis was performed only on a range of q-values, centered on the required respective (100), (012) and (002) reflection with a width of Δ*q*=0.2 nm^−1^

Following the approach described by Bernhardt et al. ^19^ the scattering distribution is treated as a probability density function for the distribution of photons. To retrieve the eigenvectors of the scattering distribution, the covariance matrix C of the distribution of the wave vector components q_y_ and q_z_ in the detection plane is diagonalized. C is defined as 

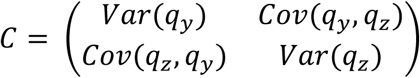

The principal direction of scattering is thereby given by the largest of the two corresponding eigenvectors v_1_ and v_2_. The length of the eigenvectors (the variance) is given by the eigenvalues λ_1_ and λ_2_. One can thereby define a dimensionless parameter, the anisotropy of the scattering: 

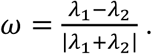

A value of 1 would thereby correspond to a single scattering direction, while a value of 0 would correspond to a perfectly isotropic scattering.

Based on our analysis, not all of crystals are aligned co-axially with respect to each other as suggested by the anisotropy of the scattering (< 0.2 for the (012) and (002) crystal planes; < 0.5 for the (100) crystal plane, see Fig. S5). The relativley low anisotropy of the scattering is probably due to the underlying disordered iridiphore layers, which reduces the levels of anisotropy. The anisotropy maps corresponding to Figs. 4A and 4D are shown in Figs. S5 C and D, respectively. Disregarding a particular reflection, one should keep in mind, that at any scan point from within the iris, at least one of the crystal planes meet the diffraction condition. This is shown in the map shown in Fig. S5 B which integrates the scattered intensity that is above a manually chosen threshold.

Several structural parameters can be extracted from a single scattering pattern. It is therefore informative to use multiple representations of the scattering data. First, in Fig. S5 A we show the scattered intensity integrated within the 10.5 nm^−1^ and 11.5 nm^−1^. The q-range was chosen based on the fact that within this range, no reflections occur. We hereby obtain a dark field contrast that is solely based on the scattering of the isotropic sample matrix and solution. In this contrast, one can for example observe trapped air bubbles in the sample preparation. The overall distribution of crystals in the sample can be obtained by integrating the scattered intensity above a manually chosen threshold that discriminates between Bragg reflections and background scattering. An example is shown in Fig. S5B. Furthermore, the three Bragg reflections can be clearly seen in maximum intensity projections of the entire dataset, see Fig. S5E. To generate a maximum intensity projection, a given pixel with index (i,j) is assigned the maximum value of all pixels with index (i,j) in the entire data set. Clearly, all reflections can hereby be visualized while in an average scattering pattern, a single reflection would be averaged out and barely visible. For comparison, a single diffraction pattern is shown in Fig. S5F.

In addition to the contrasts presented so far, one can furthermore radially integrate a single scattering pattern within a given q-range to yield a one-dimensional representation of the intensity as a function of azimuthal angle I(phi). We have performed a radial integration on the (100) reflection for each scan point. We then calculated the normalized cross-correlation of the radial intensity of each scan point with its next neighbors. We found that adjacent scan points are well correlated over a distance of 4 μm (distance between two scan points), especially, where the anisotropy was high. An overview over the entire sample as well as a zoom region is shown in Figs. S5 G and H, respectively.

## Acknowledgments

We Thank Dr. Nadav Elad for his help with the TEM Studies and Dr. Eyal Shimoni and Dr. Ifat Kaplan-Ashiri for their help with the SEM Studies. We thank Tal Bigdary for graphical schemes. We thank Prof. Leslie Leiserowitz for helpful discussions. We acknowledge the European Synchrotron Radiation Facility for provision of synchrotron radiation facilities and we would like to thank Manfred Burghammer forssistance in using beamline ID13 – microfocus beamline. We are also very thankful for helpful advice on the x-ray data analysis by Prof. Dr. Tim Salditt, Institute for X-ray Physics, Göttingen, Germany. This work was supported the Crown Center of Photonics, and the I-CORE (Israeli Center for Research Excellence) “Circle of Light”, the Israel Science Foundation (#1511/16); F.I.R.S.T. (Bikura) Individual Grant (# 2137/16); Israel Ministry of Agriculture Chief Scientist Office (#894-0194-13 and #30-04-0002); minerva-Weizmann program and Adelis Metabolic Research Fund, (in the frame of the Weizmann Institute). G.L. is an incumbent of the Elias Sourasky Professorial Chair.

## Supplementary Materials

### Supplementary Figures

**Fig. S1.**
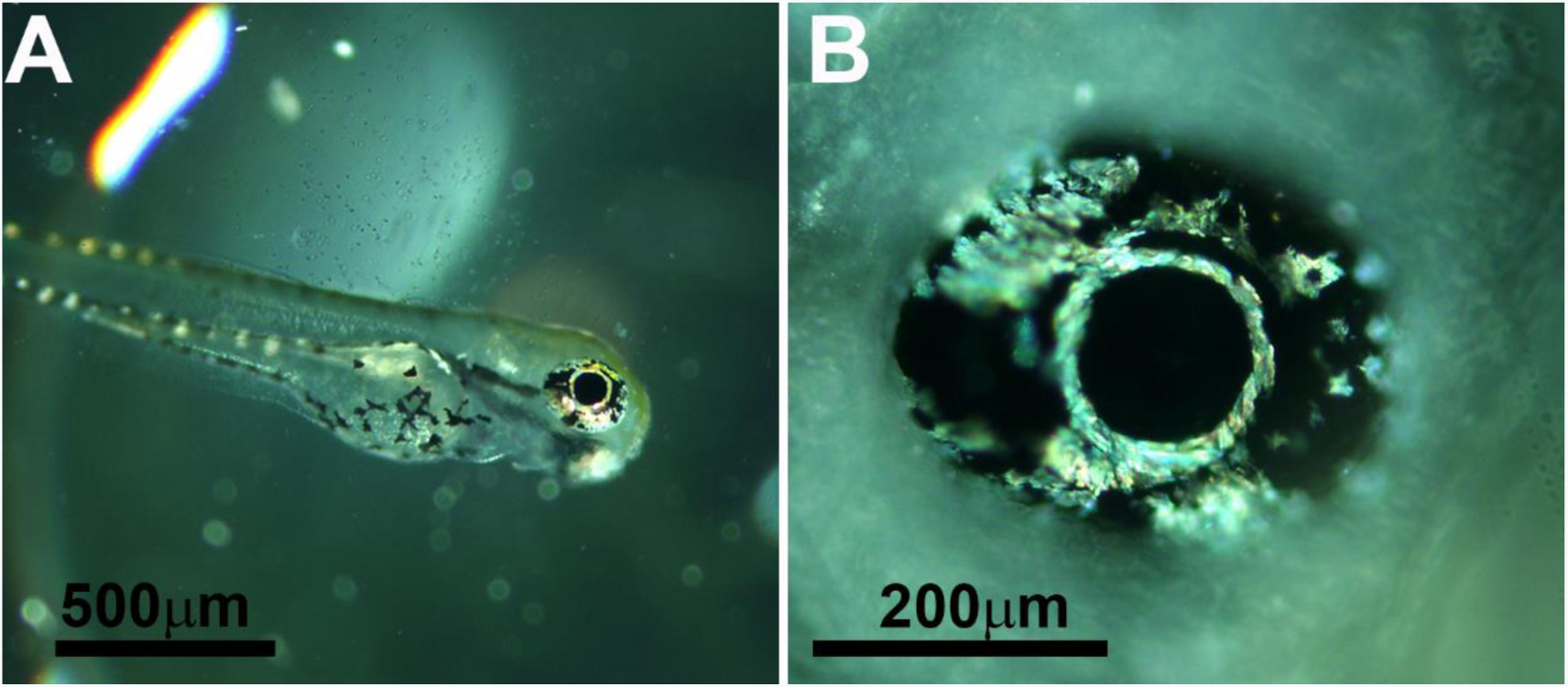
Light microscopy images of 3 dpf zebrafish larvae. (A) A complete larva. (B) Magnification of the eye. Already at this stage, the iridophores in the larval iris are clearly visible. The iridophores surrounding the lens seem to be the first to form. Scale bars: 500 μm (A), 200 μm (B),

**Fig. S2.**
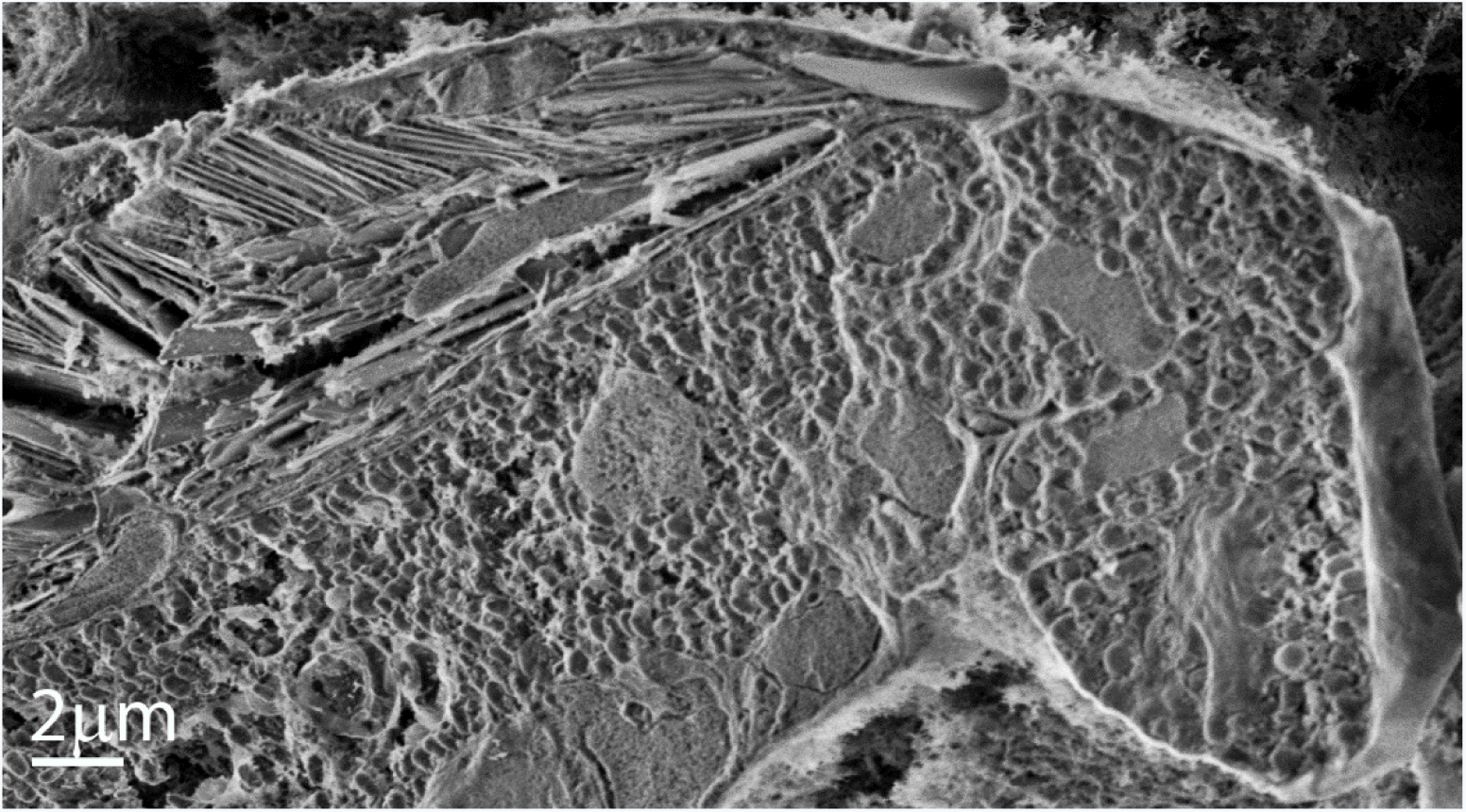
Cryo-SEM image of the zebrafish iris at 21 dpf. Image of a freeze-fractured iris shows the area adjacent to the lens, where two layers of iridophores are visible. Scale bar: 2 μm.

**Fig. S3.**
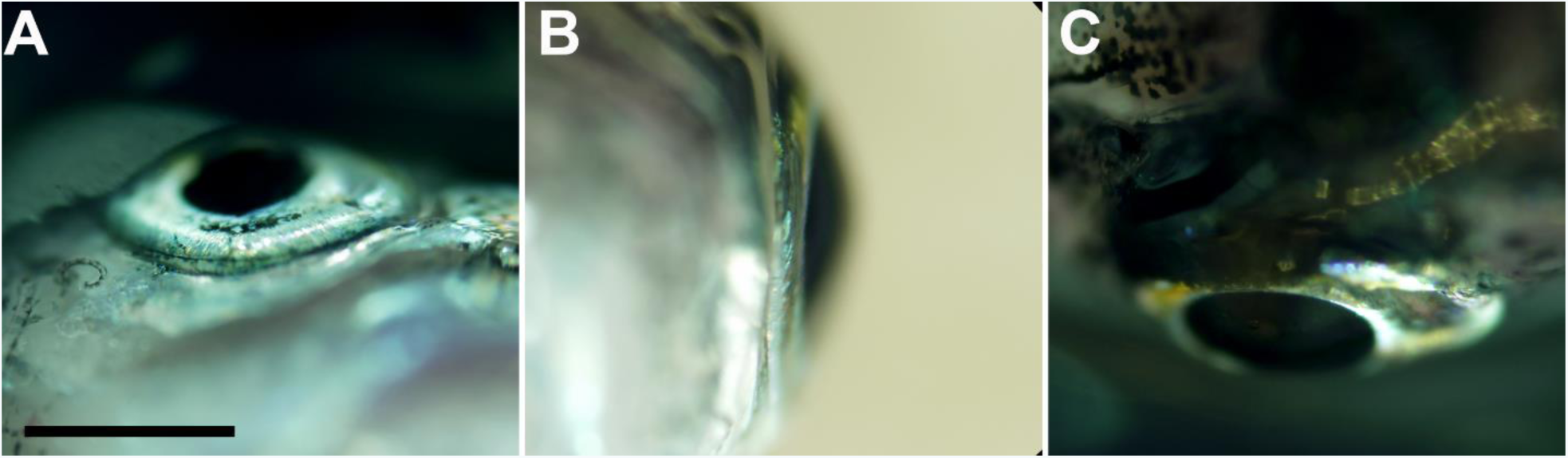
Light microscope images of the outer iridophores surrounding the eye. This layer extends to the point where the eye is protruding out from the fish head. A) Ventral view of the eye. B) A side view from posterior to anterior. C) A dorsal view of the eye.

**Fig. S4.**
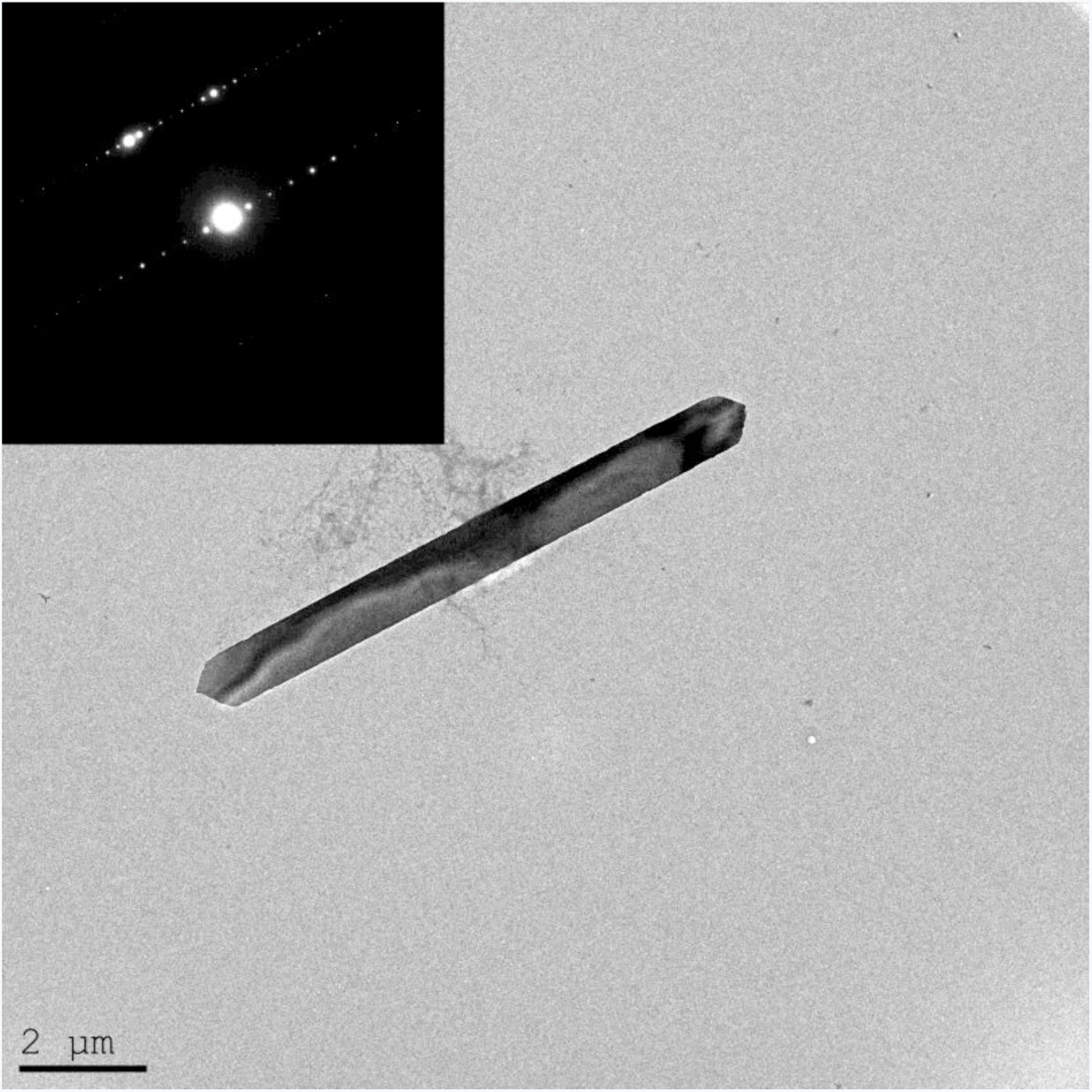
TEM images of a single guanine crystal. A TEM image showing a single crystal extracted from the zebrafish iris and its electron diffraction (inset). The electron diffraction correspond to anhydrous β-guanine.

**Fig. S5.**
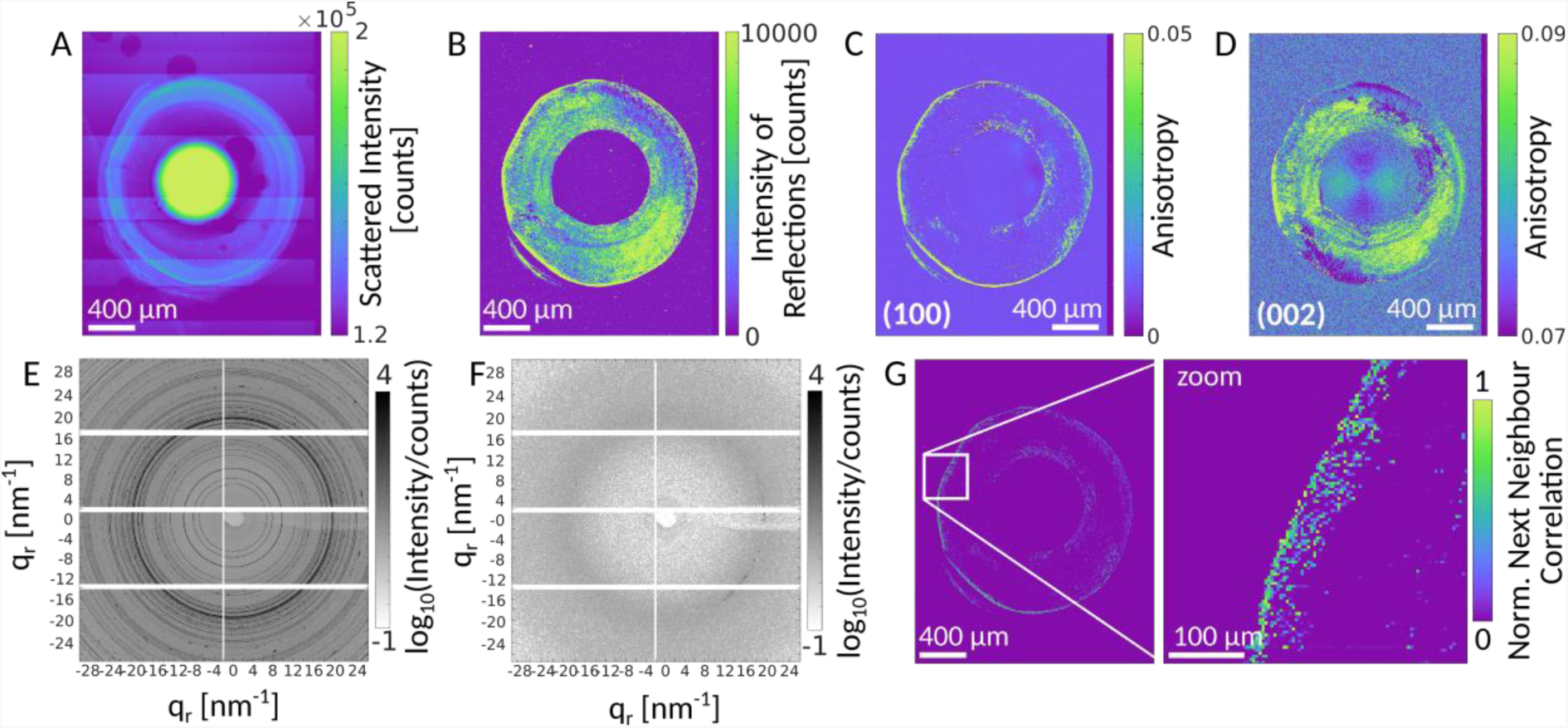
Analysis of diffraction patterns. (A) Integrated intensity between 10.5 nm^−1^ and 11.5 nm^−1^. (B) Sum over all pixels with an intensity greater than 50 counts. (C,D) Anisotropy of the (100) and (002) reflections. (E) Maximum intensity projection and (F) Isolated diffraction pattern. (G) Correlation of the radial intensity of the (100) reflection with data from its next neighbors. A zoom is shown on the right.

**Fig. S6.**
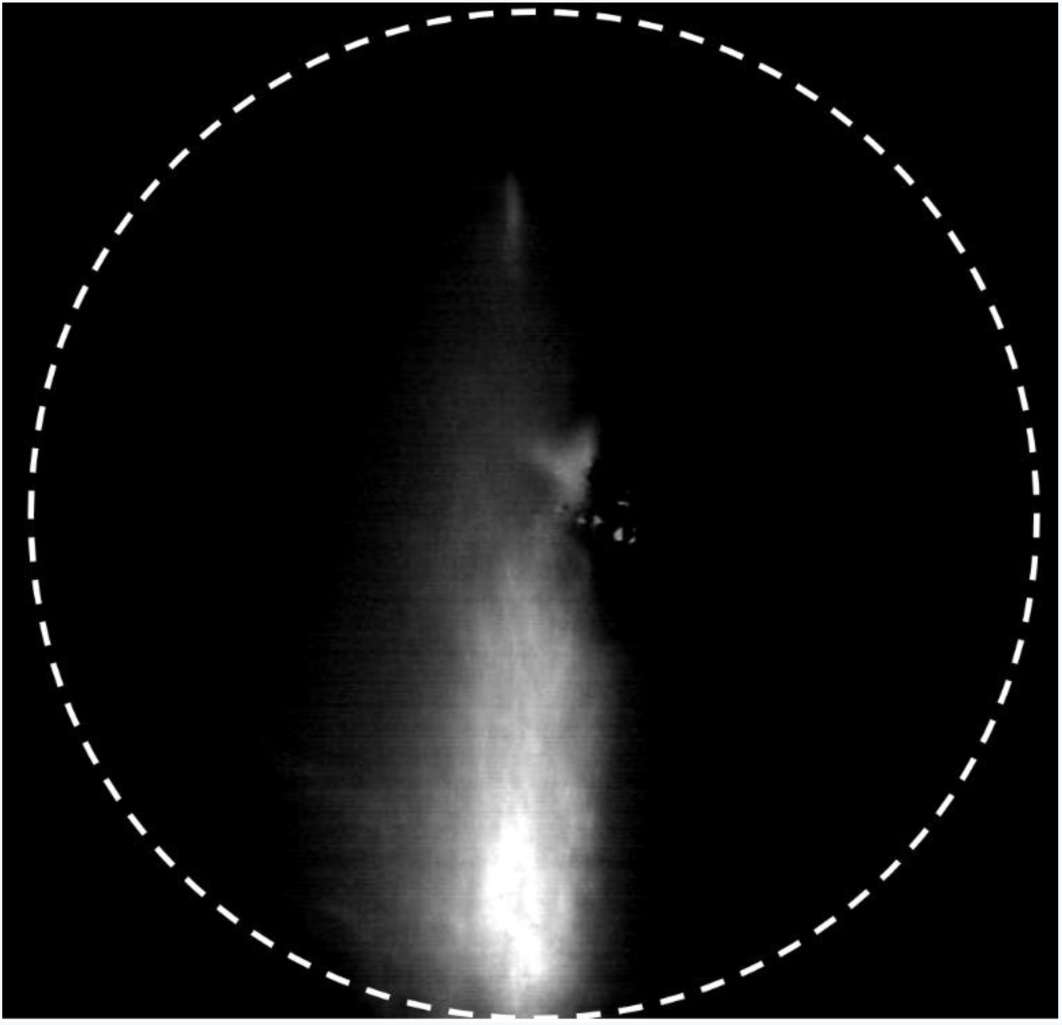
Fourier transform image of an adult zebrafish iris reflectance. A microscopic image showing the highly directional reflection of an adult zebrafish iris.

